# Unraveling the mechanisms of PAMless DNA interrogation by SpRY Cas9

**DOI:** 10.1101/2023.06.22.546082

**Authors:** Grace N. Hibshman, Jack P. K. Bravo, Hongshan Zhang, Tyler L. Dangerfield, Ilya J. Finkelstein, Kenneth A. Johnson, David W. Taylor

## Abstract

CRISPR-Cas9 is a powerful tool for genome editing, but the strict requirement for an “NGG” protospacer-adjacent motif (PAM) sequence immediately adjacent to the DNA target limits the number of editable genes. To overcome the PAM requirement, a recently developed Cas9 variant, called SpRY-Cas9 was engineered to be “PAMless” (1, 2). However, the molecular mechanisms of how SpRY can recognize all potential PAM sequences and still accurately identify DNA targets have not been investigated. Here, we combined enzyme kinetics, cryo-EM, and single-molecule imaging to determine how SpRY interrogates DNA and recognizes target sites for cleavage. Divergent PAM sequences can be accommodated through conformational flexibility within the PAM-interacting region of SpRY, which facilitates tight binding to off-target DNA sequences. Once SpRY correctly identifies a target site, nuclease activation occurs ∼1,000-fold slower than for *Streptococcus pyogenes* Cas9, enabling us to directly visualize multiple on-pathway intermediate states. Insights gained from our intermediate structures prompted rationally designed mutants with improved DNA cleavage efficiency. Our findings shed light on the molecular mechanisms of PAMless genome editing with SpRY and provide a framework for the design of future genome editing tools with improved versatility, precision, and efficiency.

## Introduction

CRISPR-Cas9 (clustered regularly interspaced short palindromic repeats, CRISPR-associated) is an RNA-guided endonuclease that uses RNA-DNA complementarity to precisely target and cleave double-stranded DNA (3). To find its target, Cas9 must meticulously search double-stranded DNA across the entire genome to identify regions of gRNA complementarity (4, 5). This search is simplified via the protospacer-adjacent motif (PAM) located immediately next to the targeted sequence (6, 7). The PAM acts as a rapid and efficient initial filter for possible target sites, circumventing the slow and energetically unfavorable process to initiate duplex melting. In the case of *Streptococcus pyogenes* Cas9 (henceforth referred to as Cas9), the required PAM sequence is NGG (where N can represent any nucleotide and G signifies guanosine). In its native context, the PAM also serves as a mechanism to distinguish between self and non-self sequences (8–10).

For genome editing applications, the PAM requirement imposes a major limitation on the number of sequences that can be targeted (2). Protein engineering efforts have focused on re-targeting or reducing the PAM specificity (11–16). A PAMless Cas9 variant, termed SpRY, is unique in that it is capable of targeting nearly any site *in vivo* (1). However, since PAM recognition is a prerequisite for efficient DNA melting and R-loop formation, it is unclear how SpRY can identify diverse target sequences.

## Promiscuous PAM recognition by SpRY

The Cas9 residues R1333 and R1335 confer specificity for two guanines in the second and third positions of the PAM by forming four hydrogen bonds with the Hoogsteen faces (8). Both residues are mutated in SpRY (R1333P, R1335Q) along with nine additional mutations (A61R, L1111R, D1135L, S1136W, G1218K, E1219Q, N1317R, A1322R, R1333P, R1335Q, and T1337R) that enable PAMless DNA targeting.

To understand how SpRY can recognize diverse PAM sequences, we determined the structures of SpRY bound to NGG, NAC, and NTC PAM-containing DNA substrates with global resolutions of 3.3, 2.8 and 3.7 Å, respectively (**Fig. 1 and Extended Data Fig. 1, Extended Data Table 1**). All three structures were in the product state, with HNH repositioned at the cleaved scissile phosphate of the target strand (TS).

**Figure 1:**
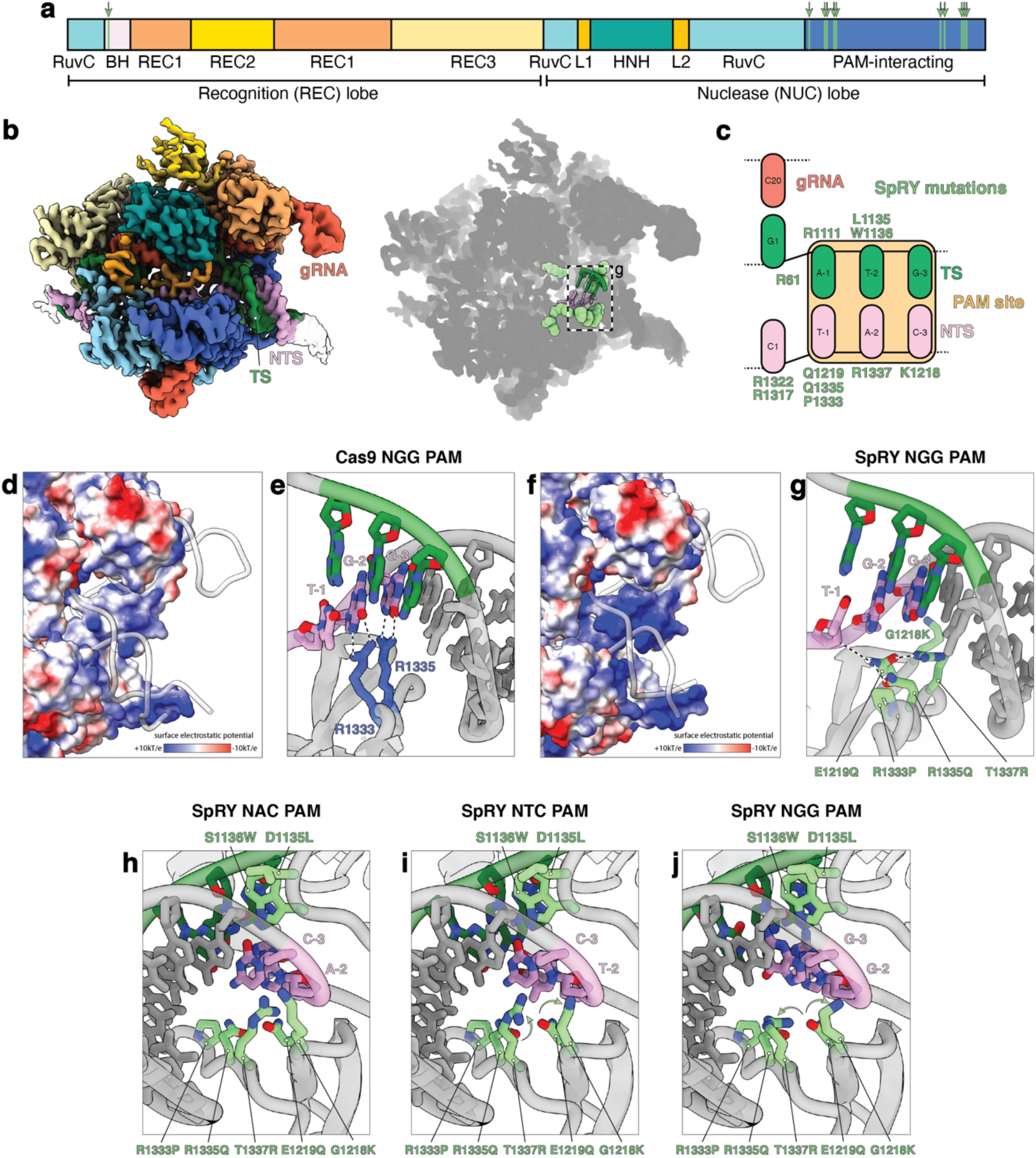
Structural basis of PAMless DNA recognition. **a**, Cas9 domain organization with SpRY mutations highlighted in light green. **b**, 2.8 Å cryo-EM reconstruction of SpRY bound to target DNA with TAC PAM sequence in the product state. Shadow of the SpRY TAC PAM product state structure with the PAM nucleotides shown as sticks and SpRY amino acid mutations shown as spheres. Molecules are colored: TS green; NTS, pink; gRNA, red; SpRY mutations, light green. **c**, Schematic representation of SpRY PAM site interaction and cleavage centers. The PAM site is highlighted in gold; HNH, dark cyan; RuvC, sky blue; Mg^2+^ in gray. **d**, Surface electrostatic potential map of Cas9. **e**, Detailed Cas9 NGG PAM site interaction (PDB: 7s4x). **f**, Surface electrostatic potential map of SpRY. **g**, Detailed SpRY NGG PAM site interaction. **h**, View of TAC, **i**, TTC, and **j**, TGG PAM site interaction turned 180° relative to **g**.

Comparison of these atomic models with a previously determined structure of Cas9 in the product state (PDB 7S4X (17)) shows that most of the complex is unaltered (RMSD of <1Å), except for the PAM-interacting (PI) domain. The mutated residues L111R, G1218K, E1219Q, R1335Q, and T1337R make several electrostatic interactions with the target DNA duplex and increase the overall net charge of the PI domain relative to Cas9 (**Fig. 1d-g**). This change in protein surface potential may stimulate the formation of non-specific electrostatic interactions with DNA to compensate for the loss of specific hydrogen bond contacts with the PAM nucleobases, enabling DNA interrogation.

Structures of the PAM-reprogrammed Cas9 variants VQR and VRQR exhibit a slight shift in the PAM region of the duplex backbone by ∼1-2 Å (18, 19). We observed a similar phenomenon in SpRY where the DNA backbone is shifted by up to 6 Å relative to the equivalent position for Cas9. This shift in the DNA backbone is mediated by the bulky, hydrophobic mutations D1135L and S1136W wedged into the minor groove.

In addition to the SpRY mutations that directly engage with the PAM, three more mutations surround the PAM readout region. N1317R and A1322R form electrostatic interactions with the phosphate backbone of the displaced NTS immediately adjacent to the PAM. These interactions likely favor initial unwinding of the DNA duplex. Additionally, A61R forms ionic interactions with E1108 at the base of the bridge helix and U63 of the gRNA scaffold, directly behind the first base of the R-loop, potentially supporting R-loop initiation (**Extended Data Fig. 2**).

**Figure 2:**
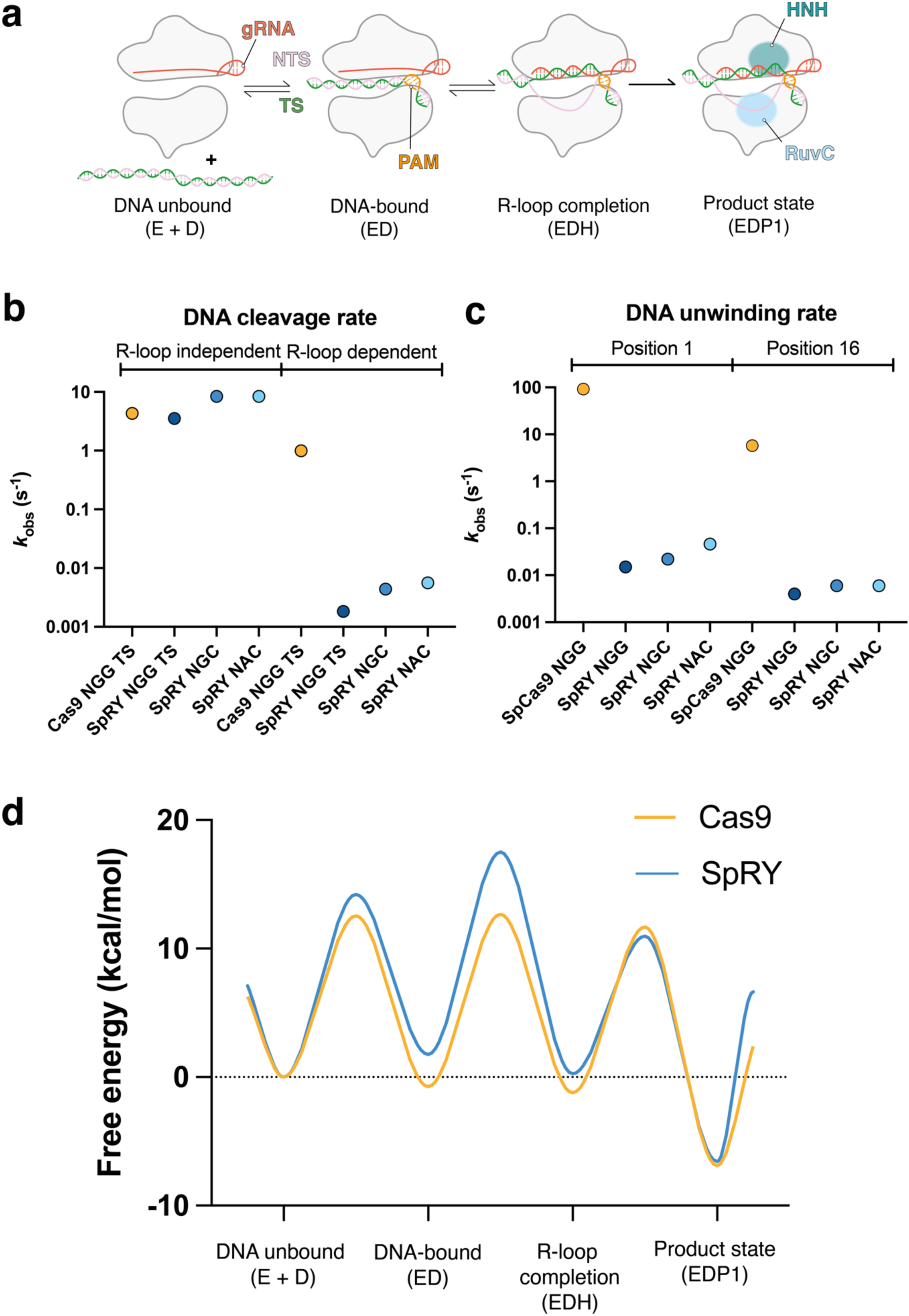
SpRY slowly unwinds DNA to find its target. **a**, Schematic representation of Cas9 enzymes reaction pathway **b**, SpRY and Cas9 observed R-loop independent (Mg^2+^-activated) and R-loop dependent (DNA-activated) cleavage rates. **c**, SpRY and Cas9 observed rates of R-loop formation at position 1 (PAM-proximal) and position 16 (PAM-distal). **d**, Free energy profile for SpRY cleavage. Model was truncated to first irreversible step (HNH cleavage of TS). Full scheme and rate constants derived from the global fitting are in **Extended Data** Fig 5, Extended Data Table 1)

Structures of SpRY in complex with NGG, NAC, and NTC PAM sequences are largely similar. Surprisingly, G1218K, N1317R, A1322R, and T1337R adopt different rotamers in the structures to maximize the energetics of binding through the formation of non-specific interactions compatible with the different PAM sequences (**Fig. 1h-j and Extended Data Fig. 2**). This is in stark contrast to Cas9 in which R1333 and R1335 are pre-ordered and poised to quickly recognize a PAM. This observation reveals a mechanism in which SpRY recognizes DNA targets through a mechanism predominantly reliant on electrostatic contacts rather than base-specific contacts.

## SpRY DNA cleavage is limited by reduced rates of DNA unwinding

Next, we analyzed DNA cleavage kinetics across different PAM sequences using the same framework we previously employed to directly compare high-fidelity variants to wild type Cas9 (20) (**Fig. 2**). Others have demonstrated essentially PAMless activity *in vitro* with assays using all possible PAM sequences for SpRY (2). We focused our efforts on NGG and two alternative PAM sequences only accessible to SpRY for more detailed kinetic analysis. In this way, we can directly compare SpRY and Cas9, and explore the efficiency of SpRY across diverse sequences.

We first measured the rate of R-loop dependent DNA cleavage by SpRY, by mixing preformed SpRY:gRNA complex with DNA in the presence of MgCl_2_. We observed that SpRY cleaved the TS at a rate of 0.0025 s^-1^, and the NTS at a rate of 0.0023 s^-1^, corresponding to a ∼500-fold slower observed rate than that of Cas9 (21). The slow observed rate of cleavage was not unique to the NGG PAM sequence, as SpRY cleaved the TS of NGC and NAC PAM substates with similarly slow rates (**Fig. 2b, Extended Data Figs. 3 & 4**).

**Figure 3:**
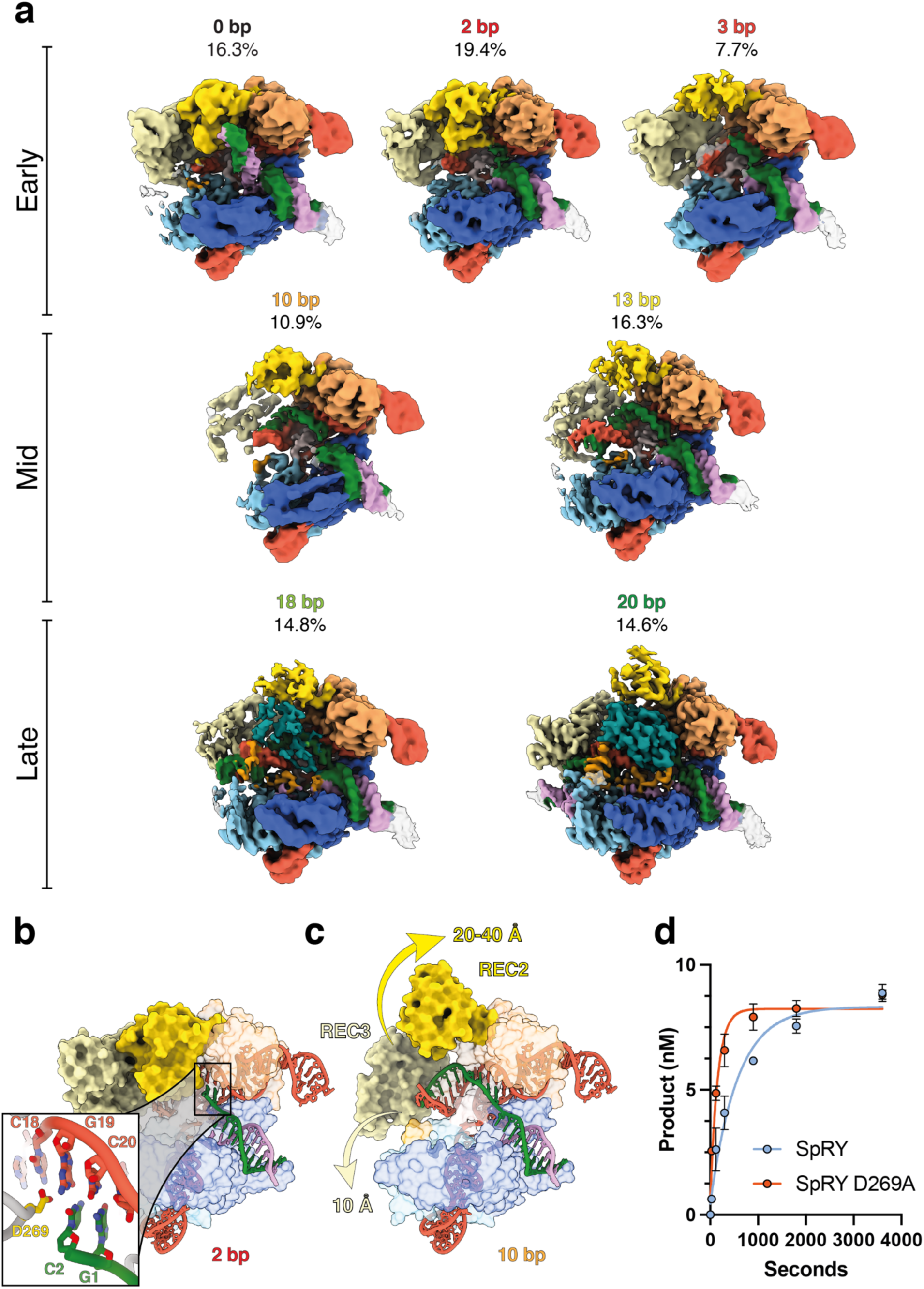
Direct visualization of SpRY R-loop formation in real-time. **a**, Ensemble cryo-EM reconstructions of SpRY during R-loop propagation with the proportion of particles as a percentage. **b & c**, Transition from early-to mid-R-loop formation requires a large conformational change of REC2 domain to create a channel to accommodate the R-loop. **d**, Alanine substitution of D269 increases the TS cleavage rate from 0.0025 s^-1^ to 0.006513 s^-1^.

**Figure 4:**
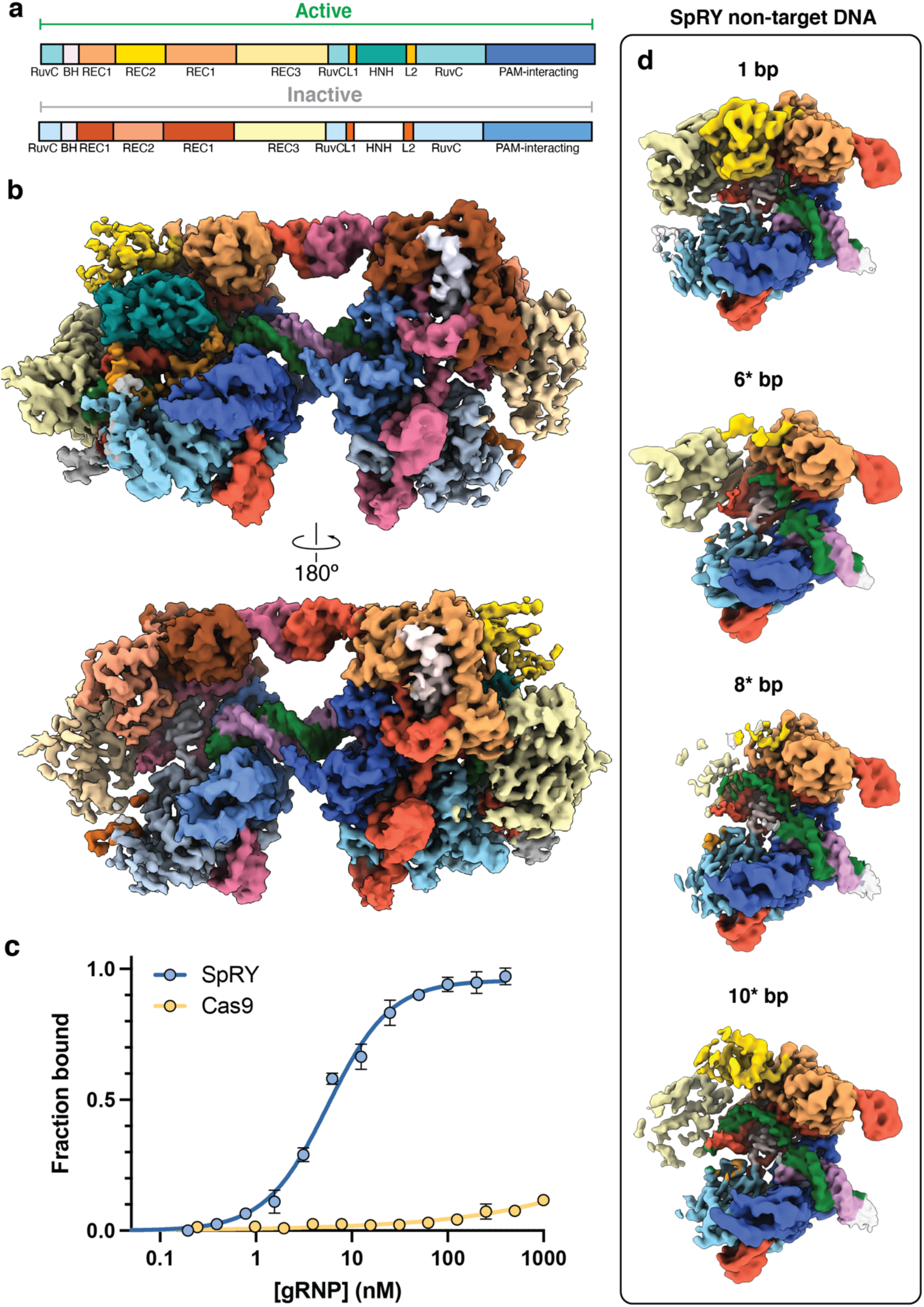
SpRY stably binds to off-target sequences. **a**, Domain organization of active (product) and inactive SpRY models in **b**. **b**, 3.0 Å cryo-EM reconstruction of the SpRYmer with one active SpRY molecule and one inactive SpRY molecule bound to the same DNA substrate. **c**, SpRY tightly binds to non-target DNA but Cas9 does not. SpRY or Cas9 were titrated into 1 nM FAM-labelled non-target DNA substrate. SpRY bound with an apparent *K*_D_ of ∼5 nM. **d**, Cryo-EM maps of SpRY-gRNA in complex with non-target DNA at different stages of non-productive R-loop formation. *6, 8, and 10 bp structures contain one, three, and five mismatches, respectively. REC2 and REC3 domains exhibit diffuse density in the 6*, 8* and 10* bp structures.

To determine whether the slow rate was due to a reduced rate of the chemical step or due to a slow step preceding chemistry, we measured R-loop independent cleavage rates by rapid quench since the reactions were too fast to resolve by hand mixing. In these experiments SpRY:gRNA was pre-incubated with DNA in the absence of magnesium and in the presence of a low concentration of EDTA to chelate any trace metal ions. The reaction was then initiated by adding MgCl_2_. We observed cleavage rates of 6 s^-1^ for the TS compared to 4.3 s^-1^ for Cas9 (21). Cleavage of the NTS for SpRY was biphasic with rates of 4.3 s^-1^ and 0.54 s^-1^ for the fast and slow phases, respectively, compared to an observed rate of 3.5 s^-1^ for Cas9 (**Fig. 2b**).

This discrepancy between R-loop dependent and independent cleavage strongly suggests that the cleavage defect is due to reduced rates of R-loop formation. To test this, we directly measured the rate of R-loop formation using a stopped-flow assay for DNA unwinding. A DNA substrate containing the fluorescent tricyclic cytosine (tC°) analog at either position 1 or position 16 on the NTS was used to report on R-loop initiation and completion. As the R-loop propagates, the NTS is displaced resulting in an increase in tC° fluorescence.

In an experiment mirroring the R-loop dependent cleavage experiment described above but performed with the fluorescent DNA substrate in the stopped flow, SpRY initiated DNA unwinding of NGG, NGC and NAC substrates with nearly identical rates (0.015, 0.022 and 0.046 s^-1^, respectively). R-loop completion was ∼3-8-fold slower (0.004, 0.006, and 0.006 s^-1^, respectively) (**Fig. 2c**). The difference between the rate of unwinding at these two positions provides direct evidence that SpRY unwinds DNA in a directional manner, from PAM-proximal to -distal, akin to Cas9. Hence, the decreased observed rate of DNA cleavage for SpRY is due to the decreased rate of DNA unwinding.

To fully understand the kinetics and mechanism of SpRY, and how it contrasts with Cas9, all the experiments for SpRY cleaving the NGG PAM substrate were fit globally to a single unified model based on numerical integration of the rate equations in KinTek Explorer (KinTek Corporation; kintekexplorer.com) (**Fig. 2d, Extended Data Fig 5, Extended Data Table 3**). We obtained a global fit to the data that could be explained by our model and a free energy profile was constructed for comparison with Cas9.. The surprising result from this analysis is that the highest barrier relative to the starting material for SpRY, defining the specificity determining step, is R-loop formation. For wild-type Cas9, barrier heights for each step were similar, suggesting that specificity is a function of all steps leading up to the chemical step.

**Figure 5:**
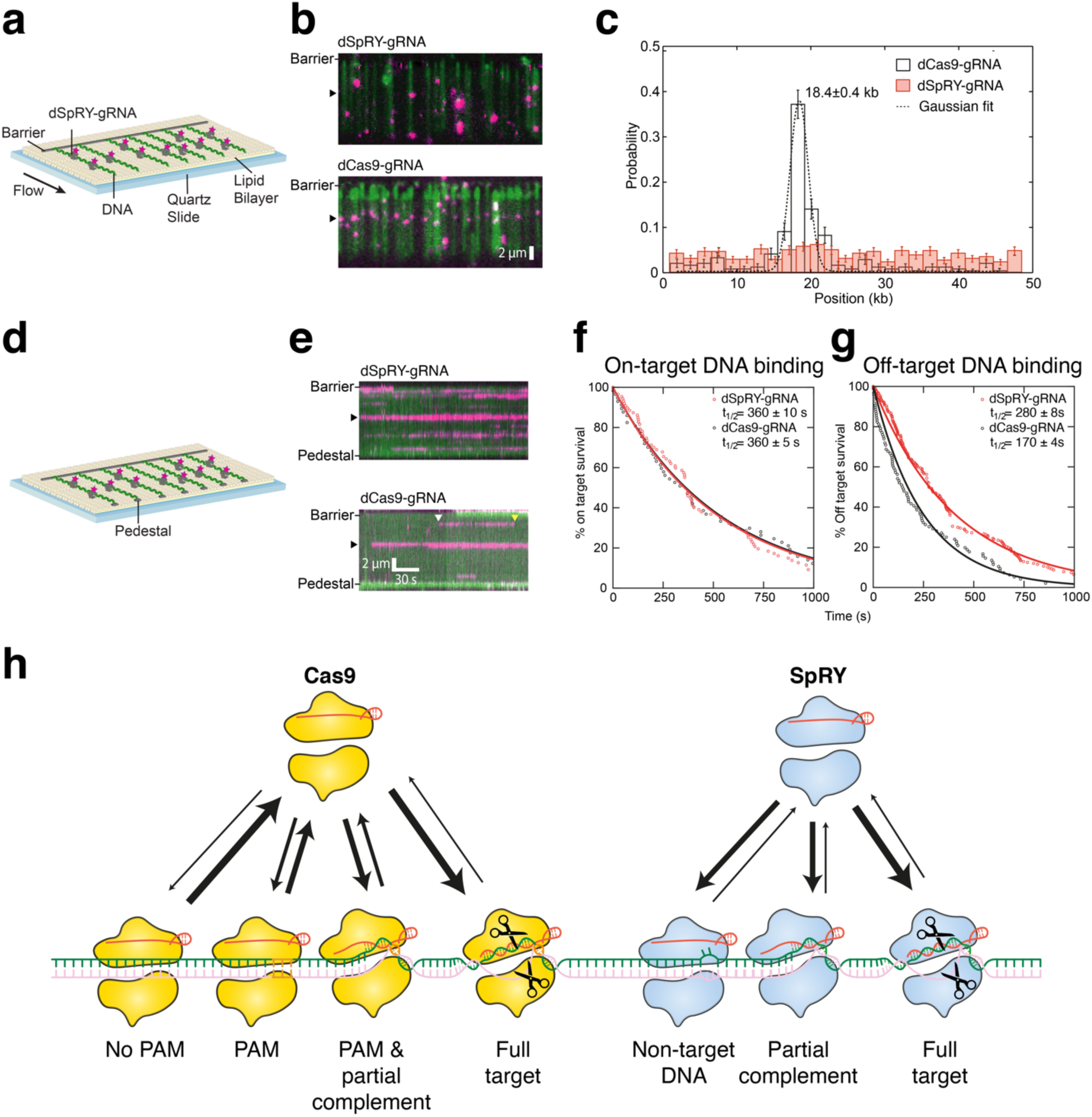
SpRY accumulates at off-target sequences during target search. **a**, Schematic of a single tether DNA curtain. **b**, YOYO-stained DNA (green) bound by fluorescently labeled dSpRY-gRNA top or dCas9-gRNA bottom (magenta). **c**, Binding distribution of dSpRY-gRNA (n=543) and dCas9-gRNA (n=242); error bars represent 95% confidence intervals. **d**, Schematic of a double tether DNA curtain. **e**, Kymographs depicting distinct binding events for dSpRY-gRNA and dCas9-gRNA. **f**, Lifetime of dSpRY-gRNA and dCas9-gRNA on target and **g,** off-target. The half-lives for each enzyme are indicated. **h**, Schematic model of target search by Cas9 and SpRY.

## Structural basis of target search and R-loop formation by SpRY

Due to the rapid rate of R-loop formation and DNA cleavage by Cas9, it has only been possible to capture intermediates of this process through covalent crosslinking, DNA substrates with limited complementarity, or catalytically dead enzyme. We reasoned that the significantly reduced enzyme kinetics may make it feasible to determine structures of SpRY during R-loop propagation without the use of non-productive substrates or chemical crosslinkers.

Contrary to typical cryo-EM sample preparation, where samples are optimized for maximal homogeneity and stability, we instead chose to prepare our sample under conditions that enrich for maximal conformational heterogeneity to trap an ensemble of on-pathway intermediate states. Based on our kinetic data, we prepared cryo-EM samples one minute after mixing SpRY:gRNA with NAC PAM DNA. At this time point, ∼90% of the DNA is unwound at position 1, less than 50% of the DNA is unwound at position 16, and only ∼10% of the DNA is cleaved. This approach captures transient states of SpRY as it initiates, propagates, and ultimately completes R-loop formation to achieve DNA cleavage. This cryo-EM dataset yielded seven high-resolution (2.9-3.4Å) structures of SpRY at distinct states along the directional R-loop propagation pathway: 0 bp, 2 bp, 3bp, 10 bp, 13 bp, 18 bp and 20 bp (product state) (**Fig. 3a and Extended Data Fig. 6**).

Since SpRY lacks a PAM requirement to bind DNA targets, it must instead identify targets through probing for sequence complementarity to the gRNA. The 0 bp structure reveals that SpRY interrogates dsDNA targets by introducing a ∼50° kink in the center of the target duplex, distorting the phosphodiester backbone immediately adjacent to the 3’ end of the gRNA spacer (**Fig. 3a**). This structure is highly similar to a recently determined structure of Cas9 covalently crosslinked to non-complementary DNA (**Extended Data Fig. 7)**. Interestingly, we did not observe an open protein/linear DNA conformation, suggesting that this conformation may be too transient to capture or a product of crosslinking.

The most populated SpRY R-loop intermediate has two bases of the TS base pairing with the gRNA (**Fig. 3a**). Within our 2 bp structure, we resolved a contact between D269 and position 3 of the gRNA spacer. This contact appears to act as a physical barrier for R-loop propagation, tethering REC2 to the seed region of the gRNA and blocking R-loop propagation (**Fig. 3b**). Transition to later R-loop intermediates is accompanied by the displacement of REC2 by ∼40Å, creating a channel to accommodate the propagating R-loop (**Fig. 3c**). Mutation of D269 to an alanine residue increased the rate of TS cleavage by ∼4-fold for the NGG PAM, confirming that disruption of this contact accelerates R-loop propagation (**Fig. 3d**). This physical barrier may be important for probing initial complementarity by making base-pairing between the gRNA and mismatch-containing duplexes unfavorable. Displacing D269 as the R-loop propagates to 3 bp unlocks REC2 from its binary position and enables further R-loop propagation (**Fig. 3a**).

The next intermediate resolved had 10 bp of R-loop formed. In this structure, the REC2 domain has been displaced, shifting upwards by up to 40 Å and REC3 is poorly resolved due to conformational flexibility. By 13 bp of R-loop formation, REC2 and REC3 are well-resolved and stabilized by R-loop contacts.

The 18 bp structure has a near-complete R-loop and exhibits partial density for the HNH domain repositioned at the scissile phosphate of the TS. When we examined the catalytic sites, we found that cleavage had not occurred at either HNH or RuvC. There is density for the required Mg^2+^ ion in the HNH active site, yet at the RuvC active site, there is only density for one of the two required Mg^2+^ ions. This structure represents a state just prior to cleavage where one of the Mg^2+^ ions is not yet properly coordinated for cleavage.

The distribution of R-loop duplex lengths we captured represent kinetic barriers for R-loop propagation. Large conformational changes are associated with each observed intermediate. While the 0 bp and 2 bp structures largely resemble the Cas9 binary complex (RMSD 1.88 Å), the transition to the 3 bp structure requires the seismic eviction of the REC2 domain. Propagation to 10 bp necessitates movement of the REC3 domain, evidenced by the lack of density in our reconstruction. By 18 bp of heteroduplex formation, HNH begins docking into the active state resulting in DNA cleavage as the R-loop completes 20 bp. Together these structures support a model of R-loop formation distinguished by structural checkpoints, as previously proposed (16, 17, 22, 23).

## SpRY tightly binds to non-target DNA sequences

During 2D classification of SpRY in complex with the NAC PAM substrate cryo-EM dataset, we observed a subset of particles that were much larger than anticipated. 3D reconstruction of these particles revealed the presence of a dimeric SpRY complex, with two SpRY gRNPs bound to the same DNA substrate – we refer to this structure as a “SpRYmer” (**Fig. 4b**). Within the SpRYmer structure, the first SpRY is in the product state, with both TS and NTS cleaved. The second SpRY resembles the binary complex, likely due to the lack of substrate complementarity necessary to initiate DNA unwinding. SpRYmer assembly is stabilized by direct linkage through the DNA and by a kissing loop interaction formed between the nexus stem loop of each gRNP (**Extended Fig. 8**). Therefore, we hypothesized that SpRY may have the capacity to stably bind to non-target DNA sequences.

To validate this, we measured the apparent binding affinity (*K*_D(app)_) of SpRY and Cas9 to an NGG PAM DNA substrate lacking complementarity to the gRNA. As anticipated, Cas9 showed no discernable binding within the concentration regime measured (up to 1000 nM) (**Fig. 4c**). This measurement agrees with previous reports where Cas9 is weakly bound to PAM sequences with an apparent affinity >10 µM (22). Strikingly, SpRY bound to this substrate with a *K*_D(app)_ of 5.6 nM, ∼2000-fold more tightly than Cas9.

To further explore this disparity in non-target DNA binding, we prepared cryo-EM samples of SpRY in complex with the same non-target DNA substrate. It became immediately clear during classification and refinement that this dataset contained a plethora of conformationally heterogeneous structures, which we separated through 3D classification. We resolved four distinct structures from this dataset, corresponding to 1 bp, 6 bp, 8 bp, and 10 bp of R-loop formation, where the 6 bp, 8 bp, and 10 bp structures contain mismatches (**Fig. 4d and Extended Data Fig. 9**).

Close inspection of the DNA sequence revealed a continuous stretch of four nucleotides with complementarity to the gRNA located within the NTS outside of the designated protospacer region. We reasoned that SpRY can associate with either DNA strand during target search, unlike Cas9 which requires the PAM sequence for proper orientation. The mismatch-containing 6, 8, and 10 bp structures align with this region of the DNA sequence where the first four bases of the R-loop are complementary to the gRNA with subsequent mismatches. Like our R-loop intermediates, we observe REC2 and REC3 movement in these structures as the R-loop propagates past that structural checkpoint.

Our structures support a model where the REC2 and REC3 domains must be displaced during R-loop formation to accommodate the propagating gRNA:TS duplex, and then repositioned to facilitate R-loop completion. The lack of PAM specificity enables SpRY to locate partial target sequences from which it cannot readily dissociate, nor achieve nuclease activation.

We propose that this high affinity binding to non-target DNA sequences is a direct consequence of the SpRY mutations which create a large, positively charged patch within the PI domain. This patch acts as an anchor for DNA binding, facilitating stable, non-specific electrostatic contacts with duplexes prior to unwinding. This novel property of SpRY compensates for the loss of low affinity, rapid PAM probing employed by Cas9, and supports dwell times long enough to search for complementarity to the gRNA.

## Single-molecule visualization of DNA interrogation by SpRY

Next, we used single-molecule DNA curtains to directly visualize DNA interrogation of dSpRY:gRNA on long DNA molecules (PMID:28668123) (**Fig. 5**). dSpRY and dCas9 were fluorescently labeled with anti-FLAG coupled quantum dots, as previously described (PMCID PMC4106473, https://doi.org/10.1101/2022.09.15.508177). We first injected Cas9 gRNP into single-tethered DNA curtains, and then flushed out excess protein with high salt buffer (**Fig. 5a-c**). As expected, dCas9 gRNPs preferentially bound to the target site (PMCID PMC4106473). In contrast, dSpRY gRNPs bound non-specifically along the entire length of the DNA with no significant enrichment at the target site (**Fig. 5a-c**).

We next used a double-tethered DNA curtain to visualize transient interactions with non-target DNA. In these assays, both DNA ends are affixed to microfabricated barriers, allowing direct observation of protein binding and release along the length of the DNA without buffer flow. This enables us to record the locations and corresponding lifetimes of all binding events (**Fig. 5d**). When bound to the target DNA sequence, the lifetime distributions for dSpRY and dCas9 were nearly identical (t_1/2_ of 360±10s and 360±5s, respectively) (**Fig. 5e & f**). In contrast, dCas9 had a relatively short binding lifetime on off-target DNA, and dSpRY remained associated with nearly the same lifetime as at the on-target sequence (**Fig 5g**). These results substantiate that the lack of a PAM requirement drastically slows target search kinetics.

## Discussion

While CRISPR-Cas9 has found widespread use as a powerful tool for programmable gene editing, the identification of target sequences by Cas9 represents a formidable challenge, akin to locating a needle within a haystack. This search predicament is simplified by the NGG PAM requirement, which significantly reduces the number of sequences that are probed for gRNA complementarity. Cas9 rapidly probes PAM sequences for an average of ∼30 ms, and far less at non-target sequences (5, 24). PAM recognition is a prerequisite for R-loop formation, which is the most significant kinetic bottleneck for DNA cleavage by Cas9 (20, 21). This process relies on the energetically unfavorable disruption of base-pairing and base stacking to expose ssDNA. Once a PAM sequence is located, the adjacent duplex is distorted to promote local base flipping and R-loop nucleation to search for gRNA complementarity (22).

Our work reveals a substantial shift in the energetic landscape governing how SpRY interacts with DNA targets differently than Cas9. As it lacks a PAM specificity, SpRY must initiate DNA melting to explore gRNA complementarity, as exemplified by our 0 bp R-loop structure (**Fig. 3a**). Considering the topological complexity of chromatin (25–27), SpRY is likely to exhibit a preference for targeting regions with low energetic barriers of DNA melting such as bends and active genes as was previously suggested (28, 29). These findings shed light on the mechanism of SpRY targeting and its propensity for specific genomic locations.

The recycling of Cas9 *in vivo* is facilitated by the recruitment of chromatin remodelers and DNA-repair factors after cleavage (5, 30–32). Unlike Cas9, SpRY stably associates with non-target sequences, as revealed by ensemble biochemistry and single-molecule imaging experiments. The accumulation of SpRY at non-target sites would result in a reduction in the effective concentration of active enzyme until it is evicted by cellular factors (33). Although this phenomenon may not drastically increase the occurrence of non-target double-strand breaks due to insufficient gRNA complementarity necessary for HNH realignment and DNA cleavage (17), it does have the potential to induce undesirable gene silencing. In comparison, Cas9 requires a PAM and at least five nucleotides of seed complementarity to stably associate with non-target sequences. This stable association of Cas9 with non-target sequences has been demonstrated to cause cellular toxicity (28, 34). These observations emphasize the importance of carefully considering the implications of SpRY and Cas9 behavior to mitigate off-target effects and enhance the safety of gene editing applications.

In conclusion, despite its limitations in cleavage efficiency, SpRY proves to be an asset in modifying genomic regions that were previously inaccessible to Cas9. Recent successful gene editing in stem cells, plants and model organisms exhibits its effectiveness (35–37). Through our structural analysis, we identified a mutation that accelerates gRNA:TS duplex formation and increases DNA cleavage efficiency (**Fig 3**). Combining this mutation with other hyperactive Cas9 mutations holds the potential to compensate for SpRY’s reduced activity (38). The creation of chimeric CRISPR-Cas9 variants, where mutants with diverse purposes are combined, could forge the development of highly specific and adaptable gene editing tools that transcend biological constraints. Our discoveries will serve as a valuable foundation for the generation of potent chimeric Cas9 variants advancing the field toward the most effective gene editing tool.

## Acknowledgements

This work was supported in part by Welch Foundation grants F-1604 (to K.A.J.), F-1938 (to D.W.T.), and F-1808 (to I.J.F), a Robert J. Kleberg, Jr. and Helen C. Kleberg Foundation Medical Research Grant (to D.W.T.), and the National Institutes of Health R01GM124141 (to I.J.F.) and R35GM138348 (to D.W.T.). The content is solely the responsibility of the authors and does not necessarily represent the official views of the National Institutes of Health. D.W.T is a CPRIT Scholar supported by the Cancer Prevention and Research Institute of Texas (RR160088). We thank I. Stohkendl in the Taylor group for insightful discussions.

## Author Contributions

Conceptualization: J.P.K.B. Investigation and data analysis: G.N.H. (protein production, cryo-EM data collection, structure determination and modelling), J.P.K.B (biochemistry and kinetic analysis), H.Z. (single-molecule imaging), T.L.D. (kinetic analysis). Writing initial draft: J.P.K.B. and G.N.H. All authors reviewed, edited and approved the manuscript. Supervision: J.P.K.B. and D.W.T. Funding acquisition: D.W.T., K.A.J., and I.J.F.

## Data Availability

The structures and associated atomic coordinates have been deposited into the Electron Microscopy Data Bank (EMDB) and Protein Data Bank (PDB) with accession codes SpRY NAC PAM 20 bp (EMD-40740 and PDB 8SRS), SpRY NGG PAM (EMD-40681 and PDB 8SPQ), SpRY NTC PAM (EMD-40705 and PDB 8SQH), SpRY NAC PAM 0 bp (EMD-41073 and PDB 8T6O), SpRY NAC PAM 2 bp (EMD-41074 and PDB 8T6P), SpRY NAC PAM 3 bp (EMD-41085 and PDB 8T76), SpRY NAC PAM 10 bp (EMD-41079 and PDB 8T6S), SpRY NAC PAM 13 bp (EMD-41080 and PDB 8T6T), SpRY NAC PAM 18 bp (EMD-41083 and PDB 8T6X), SpRY non-target 1 bp (EMD-41084 and PDB 8T6Y), SpRY non-target 6* bp (EMD-41086 and PDB 8T77), SpRY non-target 8* bp (EMD-41087 and PDB 8T78), SpRY non-target 10* bp (EMD-41088 and PDB 8T79), and SpRYmer (EMD-41093 and PDB 8T7S).

## Methods

### Protein expression and purification

Cas9 was expressed and purified as described previously (39). SpRY was expressed from pET28-SpRY-NLS-6xHis (gbr2101) purchased from Addgene (Addgene plasmid # 181743; http://n2t.net/addgene:181743 ; RRID:Addgene_181743) (2). SpRY was then expressed and purified in the same manner as Cas9.

### Nucleic acid preparation

55-nt DNA duplexes were prepared from PAGE-purified oligonucleotides synthesized by Integrated DNA Technologies. DNA duplexes used in cleavage assays were prepared by mixing 6-FAM or Cy3 labeled target strands with unlabeled non-target strands at a 1:1.15 molar ratio in annealing buffer (10 mM Tris-HCl pH 8, 50 mM NaCl, 1 mM EDTA), heating to 95°C for 5 min, then cooling to room temperature over the course of 1 hr. The sgRNA was purchased from Synthego and annealed in annealing buffer using the same protocol as for the duplex DNA substrates. The sequences of the synthesized oligonucleotide are listed in **Extended Data Table 1**.

## Kinetics

### Buffer composition for kinetic reactions

Cleavage reactions were performed in 1X cleavage buffer (20 mM Tris-Cl, pH 7.5, 100 mM KCl, 5% glycerol, 1 mM DTT) at 37 °C.

### DNA cleavage kinetics

R-loop independent DNA cleavage: The reaction of Cas9 with on-and off-target DNA was performed by preincubating Cas9.gRNA (28 nM active-site concentration of Cas9, 100 nM gRNA) with 10 nM DNA with a 6-FAM label on the target strand in the absence of Mg^2+^, and trace amounts of EDTA to chelate any co-purified cations (0.2 mM). The reaction was initiated by adding Mg^2+^ to 10 mM, then stopped at various times by mixing with 0.3 M EDTA (Extended Data Fig. 1). R-loop dependent DNA cleavage: as above, but Cas9, gRNA and Mg^2+^ were co-incubated, and the reaction was initiated by the addition of 10 nM DNA. Products of the reaction were resolved and quantified using an Applied Biosystems DNA sequencer (ABI 3130xl) equipped with a 36 cm capillary array and nanoPOP6 polymer (MCLab) (40). Data fit to equations were fit using either a single or double-exponential equations shown below:

Single exponetial equation:

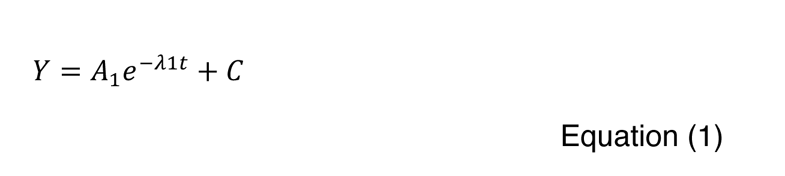

where *Y* represents concentration of cleavage product, *A_1_* represents the amplitude, and *λ*_1_ represents the observed decay rate (eigenvalue). The half-life was calculated as t_1/2_ = ln(2)/ ²A_1_.

Double exponetial equation:

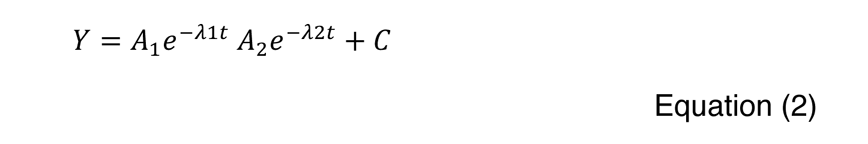

where *Y* represents concentration of cleavage product, *A_1_* represents the amplitude and *λ*_1_ represents the observed rate for the first phase. *A_2_* represents the amplitude and *λ*_2_ represents the observed rate for the second phase.

### Stopped-flow kinetic assay

Stopped-flow experiment was performed as previously described (20). Briefly, 250 nM Cas9-gRNA complex (1:1 ratio, active site Cas9 concentration) was mixed with 100 nM tC°-labeled 55/55 nt DNA substrate at 37°C using AutoSF-120 stopped-flow instrument (KinTek Corporation, Austin, TX). Excitation was at 367 nm, and emission was monitored with a 445 nm filter with a 20 nm bandpass (Semrock).

## CryoEM sample preparation, data collection, and processing

SpRY bound to different DNA substrates were assembled by mixing SpRY with gRNA in a 1:1.5 molar ratio and incubated at room temperature for 10 min in reaction buffer (20 mM Tris-Cl, pH 7.5, 100 mM KCl, 10 mM MgCl_2_, 5% glycerol, and 5 mM DTT). Each DNA substrate was then added in a 1:1 molar ratio with 10 µM SpRY gRNP. The product state PAM complexes were incubated for 1 hr at room temperature and the non-target DNA complex was incubated for 2 hr at room temperature. Based on our kinetic analysis, R-loop intermediates were prepared by triggering the DNA cleavage reaction using NAC PAM DNA and incubating with SpRY gRNP at 37°C for 60 s. The reactions were quenched by vitrification. 2.5 µl of sample was applied to glow discharged holey carbon grids (Quantifoil 1.2/1.3), blotted for 6 s with a blot force of 0, and rapidly plunged into liquid nitrogen-cooled ethane using an FEI Vitrobot MarkIV.

All datasets apart from NTC PAM were collected on an FEI Titan Krios cryo-electron microscope equipped with a K3 Summit direct electron detector (Gatan, Pleasanton, CA). Images were recorded with SerialEM(41) with a pixel size of 0.83 Å. Movies were recorded at 13.3 electrons/pixel/second for 6 s (80 frames) to give a total dose of 80 electrons/pixel. The NTC dataset was collected on a FEI Glacios cryo-TEM equipped with a Falcon 4 detector with a pixel size of 0.94 Å, and a total exposure time of 15s resulting in a total accumulated dose of 40 e/Å^2^ which was split into 60 EER fractions. All datasets were collected with a defocus range of -1.5 to -2.5 µm. Motion correction, CTF estimation and particle picking was performed on-the-fly using cryoSPARC Live v4.0.0-privatebeta.2(42). Further data processing was performed with cryoSPARC v.3.2. A total of 2,016 movies were collected for the NTC dataset, 4,293 movies for the NGG dataset, 3,546 movies for the NAC dataset, 10,151 movies for the R-loop intermediate dataset, and 8,765 movies for the non-target DNA dataset.

The NTC, NGG, and NAC datasets were processed using similar workflows starting with blob picker with a minimum particle diameter set to 100 Å, and a maximum particle diameter set to 180 Å. The particles were then subjected to a single round of 2D classification. The particles selected from 2D classification were processed via ab initio reconstruction, followed by heterogeneous refinement. After multiple rounds of ab initio reconstruction and heterogeneous refinement, the final reconstructions were formed from non-uniform refinement. There appeared to be a second SpRY molecule bound to the same DNA in some of the NAC PAM 2D classes. Particles within these classes were extracted with a larger box size of 512 pixels yielding a dimer structure. Masks for each half of the dimer were created and particle subtraction was performed on each half. The subtracted particles were then subjected to local refinement, and ultimately recombined in ChimeraX producing the SpRYmer structure.

The R-loop intermediate and non-target DNA datasets were processed following similar workflows, again starting with blob picker as in the previous datasets. After the initial round of 2D classification, ab initio reconstruction, and heterogeneous refinement, the particles were processed using 3D classification with the parameters of force hard classification and PCA initialization mode. This method generated classes with a variable number of R-loop base pairs formed. The particles corresponding to a single state were extracted with a 384-pixel box size, globally and locally CTF refined, and fed to non-uniform refinement which yielded the final reconstructions.

## Structural model building and refinement

Product state Cas9 (PDB 7S4X) was used as a starting model for the NGG PAM structure. Once built, the NGG PAM structure was used as a starting model for the NAC, NTC, 10 bp, and 18 bp structures. The 18 bp structure was used as a starting model for the 13 bp structure. SpCas9 with 0 bp, closed-protein/bent-DNA conformation (PDB 7S36) was used as a starting model for the 0 bp intermediate. SpCas9 with 3 bp R-loop (PDB 7S38) was used as a starting model for the 2 bp structure, which was then used as the starting model for the 3 bp R-loop intermediate and 1 bp non-target structure. The 3 bp R-loop intermediate was used as a starting model for the 6*bp non-target structure, which was then used as the starting model for the 8*bp non-target structure. The 10 bp R-loop intermediate was used as the starting model for the 10*bp non-target structure. SpCas9 binary complex (PDB 4ZT0) was used as the starting model for the inactive half of the SpRYmer, and the NAC PAM structure created for this manuscript was used as the starting model for the active half. Nucleic acid alterations were made in Coot*, and further modeling was performed using Isolde(43). The models were ultimately subjected to real-space refinement implemented in Phenix.

All structural figures and videos were generated using ChimeraX v1.2 (44).

## Single-molecule fluorescence microscopy

Single-molecule fluorescent images were collected using a customized prism TIRF microscopy-based inverted Nikon Ti-E microscope system equipped with a motorized stage (Prio ProScan II H117). The sample was illuminated with a 488 nm laser (Coherent Sapphire) through a quartz prim (Tower Optical Co.). For imaging SYTOX Orange-stained DNA and Anti-FLAG-Qdot705-labled dSpRY or dSpCas9, the 488 nm laser power was adjusted to deliver low power (4 mW) at the front face of the prism using a neutral density filter set (Thorlabs). The imaging was recorded using electron-multiplying charge-coupled device (EMCCD) cameras (Andor iXon DU897). Flowcells used for single-molecule DNA experiments were prepared as previously described (PMDI:32621611). Briefly, a 4-mm-wide, 100-μm-high flow channel was constructed between a glass coverslip (VWR 48393 059) and a custom-made quartz microscope slide using two-sided tape (3M 665). Double-tethered DNA curtains were prepared with 40 μL of liposome stock solution (97.7% DOPC (Avanti #850375P), 2.0% DOPE-mPEG2k (Avanti #880130P), and 0.3% DOPE-biotin (Avanti #870273P) in 960 μL Lipids Buffer (10 mM Tris-HCl, pH 8, 100 mM NaCl) incubated in the flowcell for 30 minutes. Then, 50 μg μL-1 of goat anti-rabbit polyclonal antibody (ICL Labs, #GGHL-15A) diluted in Lipids Buffer was incubated in the flowcell for 10 minutes. The flowcell was washed with BSA Buffer (40 mM Tris–HCl, pH 8, 2 mM MgCl2, 1 mM DTT, 0.2 mg mL-1 BSA) and 1 μg L-1 of digoxigenin monoclonal antibody (Life Technologies, #700772) diluted in BSA Buffer was injected and incubated for 10 minutes. Streptavidin (0.1 mg mL-1 diluted in BSA Buffer) was injected into the flowcell for another 10 minutes. Finally, ∼12.5 ng μL-1 of DNA substrate was injected into the flowcell. To prepare single-tethered DNA curtains, the anti-rabbit antibody and digoxigenin antibody steps were omitted.

For single-tethered DNA experiments, DNA was visualized in the presence of a continuous flow (0.2 mL min^-1^) in the imaging buffer (40 mM Tris-HCl pH 8.0, 2 mM MgCl2, 0.2 mg mL^-1^ BSA, 50 mM NaCl, 1 mM DTT, 100 nM SYTOX Orange) supplemented with an oxygen scavenging system (3% D-glucose (w/v), 1 mM Trolox, 1500 units catalase, 250 units glucose oxidase). For double-tethered DNA experiments, the flow was turned off after protein entered into flowcell. NIS-Elements software (Nikon) was used to collect the images at 500 ms frame rate with 100 ms exposure time. All images were exported as uncompressed TIFF stacks for further analysis in FIJI (NIH) and MATLAB (The MathWorks). The half lifetime of dSpRY and dSpCas9 were fitted by single-exponential equation.

## DNA substate preparation for single-molecule assays

To prepare DNA substates for microscopy, 125 μg of λ-phage DNA was mixed with two oligos (2 μM oligo Lab07 and 2 μM oligo Lab09) in 1× T4 DNA ligase reaction buffer (NEB B0202S) and heated to 70°C for 15 min followed by gradual cooling to 15°C for 2 hours. One oligo will be annealed with the overhand located at the left cohesive end of DNA, and the other oligo will be annealed with the overhand at right cohesive end. After the oligomer hybridization, 2 μL of T4 DNA ligase (NEB M0202S) was added to the mixture and incubated overnight at room temperature to seal nicks on DNA. The ligase was inactivated with 2 M NaCl, and the reaction was injected to an S-1000 gel filtration column (GE) to remove excess oligonucleotides and proteins.

## dSpRY-gRNA and dSpCas9-gRNA preparation and fluorescent labeling

dSpRY and dSpCas9 ribonucleoprotein complexes were reconstituted by incubating a 1:2 molar ratio of apoprotein and sgRNA (sequence: GUG AUA AGU GGA AUG CCA UGG UUU UAG AGC UAG AAA UAG CAA GUU AAA AUA AGG CUA GUC CGU UAU CAA CUU GAA AAA GUG GCA CCG AGU CGG UGC UUU U, λ target-18.1 kb) in incubation buffer (50 mM Tris-HCl pH 8.0, 5 mM MgCl_2_, 2 mM DTT) followed by incubation at 25 °C for 25 minutes. dSpRY and dSpCas9 were labeled by anti-FLAG antibody coupled with Qdot705, and then diluted to 0.5 nM – 2 nM in imaging buffer before injected into the flowcell. For dSpRY-gRNA and dSpCas9-gRNA binding experiments, single-tethered DNA curtains were assembled and protein was diluted and incubated with sgRNA as described above. Ribonucleoprotein was diluted in imaging buffer to a final concentration of 2 nM. For dSpRY-gRNA and dSpCas9-gRNA binding lifetime experiments, double-tethered DNA curtains were assembled and ribonucleoprotein was diluted in imaging buffer to a final concentration of 0.5 nM.

## Fluorescence anisotropy binding assays

SpRY and Cas9 gRNP complexes were prepared as described above, and serial 2-fold dilutions were incubated with 0.8 nM FAM-labelled scrambled DNA for 3h at 37°C in 1 x Cas9 buffer supplemented with 0.05 % Tween-20. Data were recorded at 37°C in a CLARIOstar Plus multi-detection plate reader (BMG Labtech) equipped with a fluorescence polarization optical module (*λ*_ex_ = 485 nm; *λ*_em_ = 520 nm). The data were normalised and binding curves were fitted in Graphpad Prism 9 using a hyperbola.

## Extended Data Figures & Tables

**Extended data figure 1.**
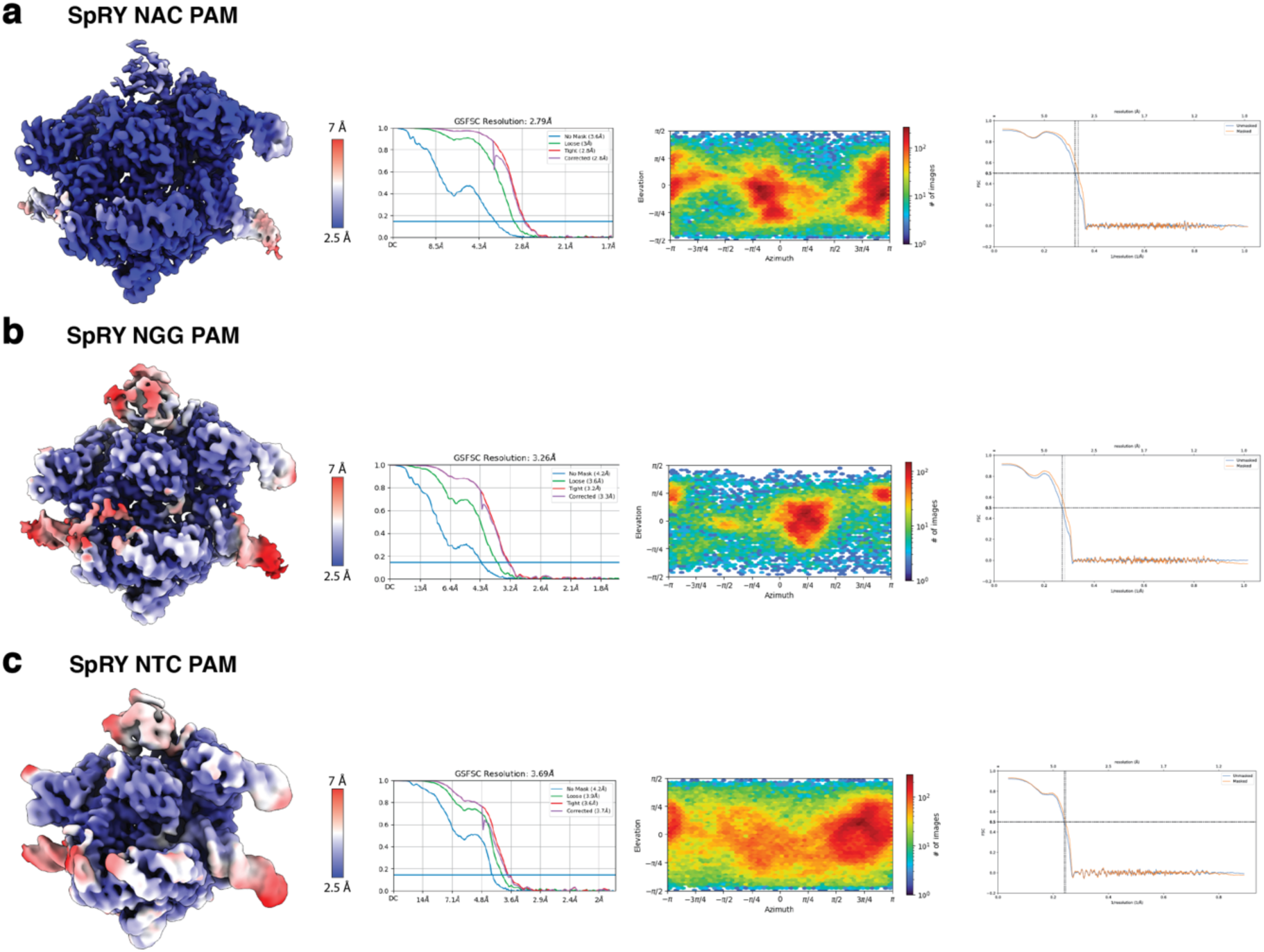
Cryo-EM data analysis of SpRY **a**, NAC PAM DNA **b**, NGG PAM DNA, and **c**, NTC PAM DNA.

**Extended data figure 2.**
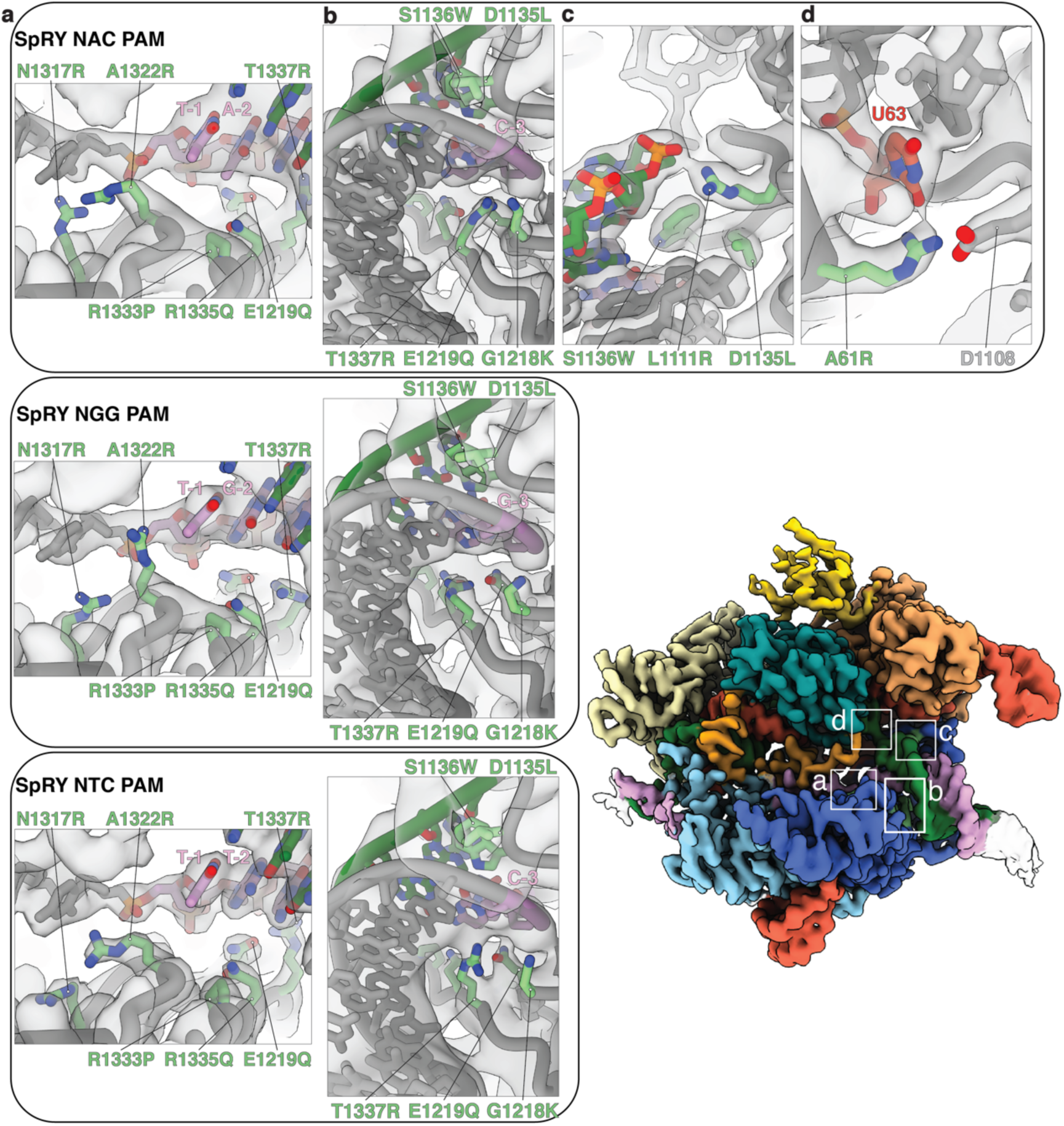
Detailed view of SpRY mutations and the rotamers adopted by different PAM sequences.

**Extended data figure 3.**
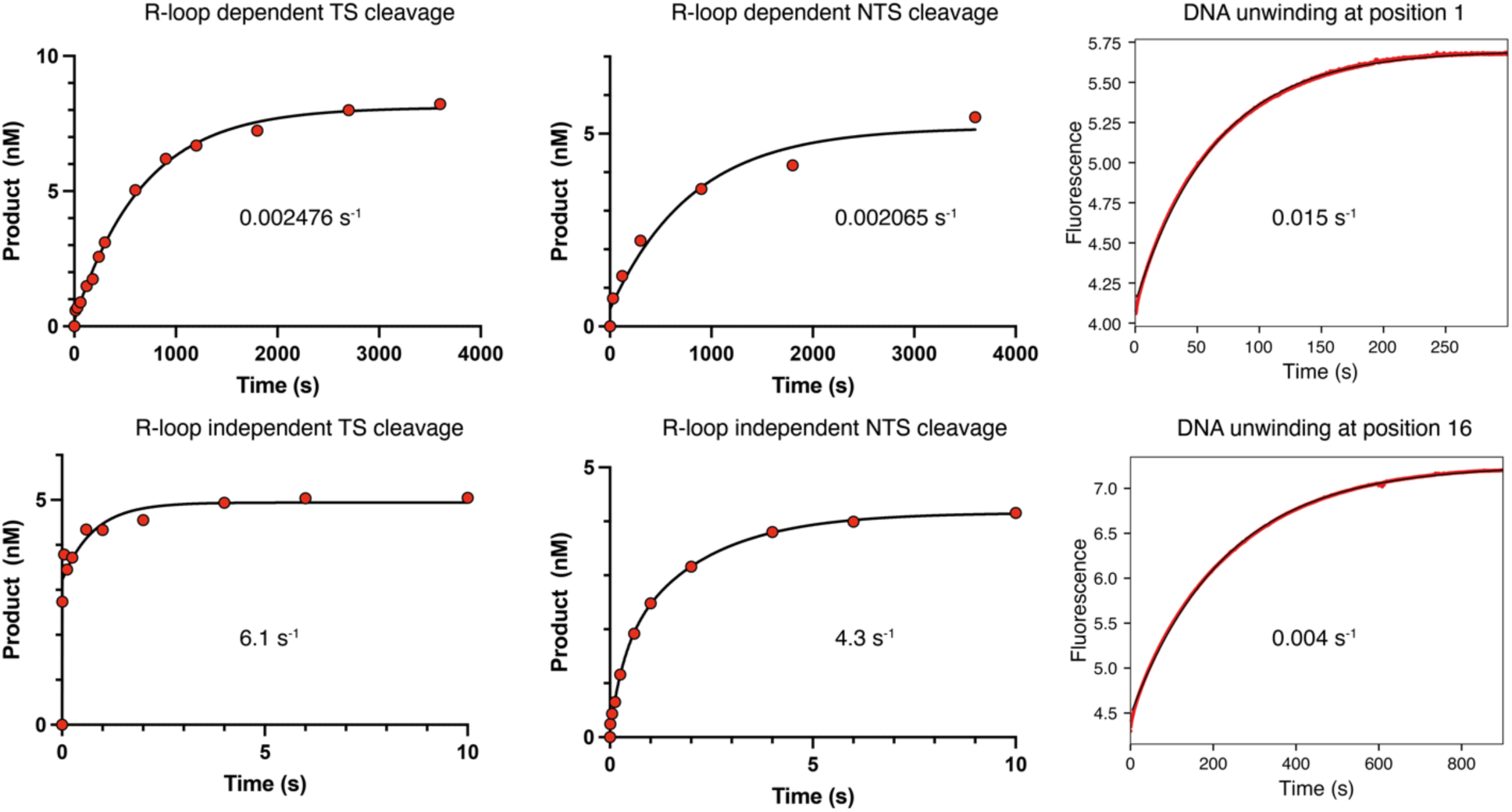
NGG PAM DNA cleavage kinetics.

**Extended data figure 4.**
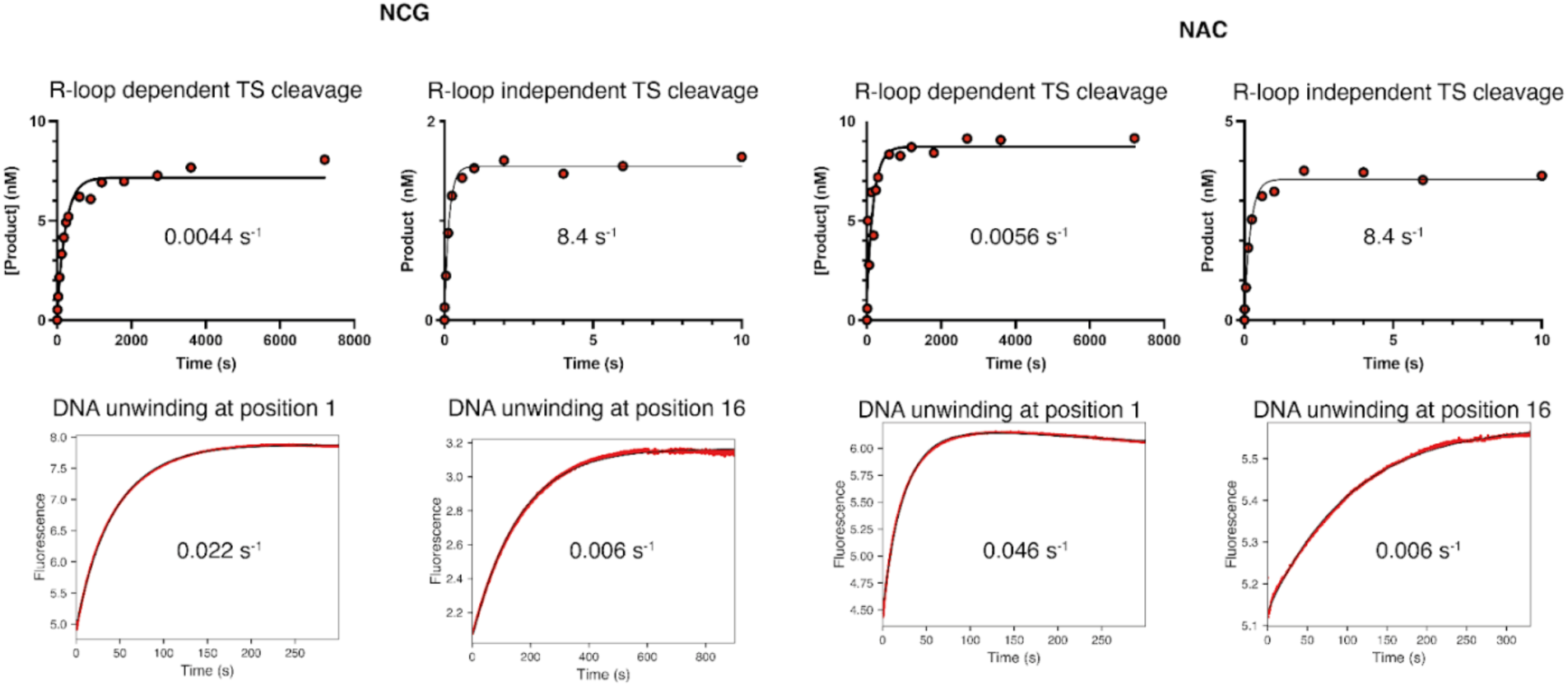
DNA cleavage and unwinding kinetics for NGC and NAC PAM substrates. The low amplitude of product for the Mg^2+^-initiated reactions may be due to association of SpRY at partially complementary, non-target sites, thus reducing the effective enzyme concentration.

**Extended data Figure 5.**
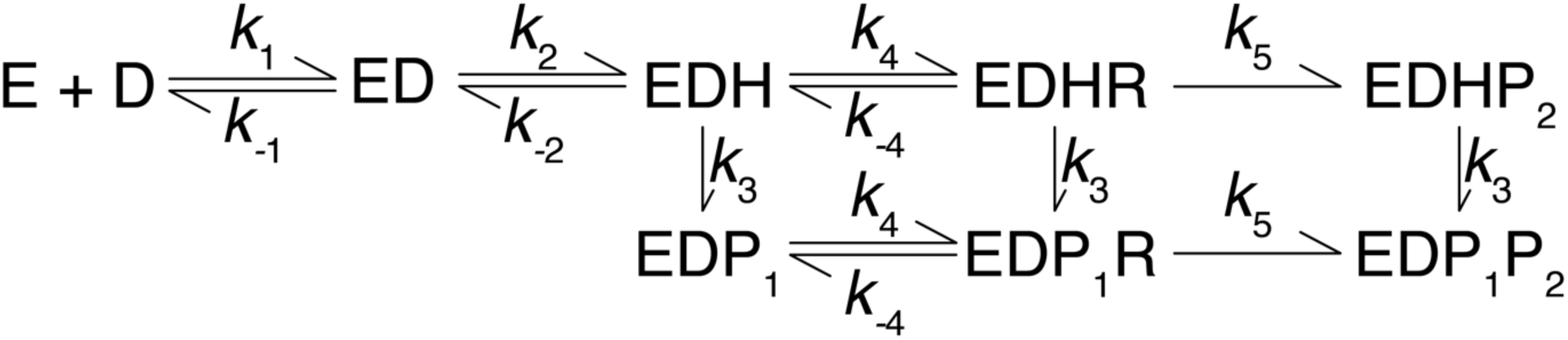
E is Cas9.gRNA, D is target DNA, ED is Cas9.gRNA.DNA, EDH is R-loop formation with docking of target strand to HNH, EDHR and EDP1R are R-loop formation with docking of non-target strand to RuvC, EDHP2 is RuvC cleavage of non-target strand, EDP_1_ is HNH cleavage of the target-strand, EDP_1_P_2_ is cleavage of both strands, Rates obtained from Fitting in KinTek Explorer are shown in Extended data table 1.

**Extended data figure 6.**
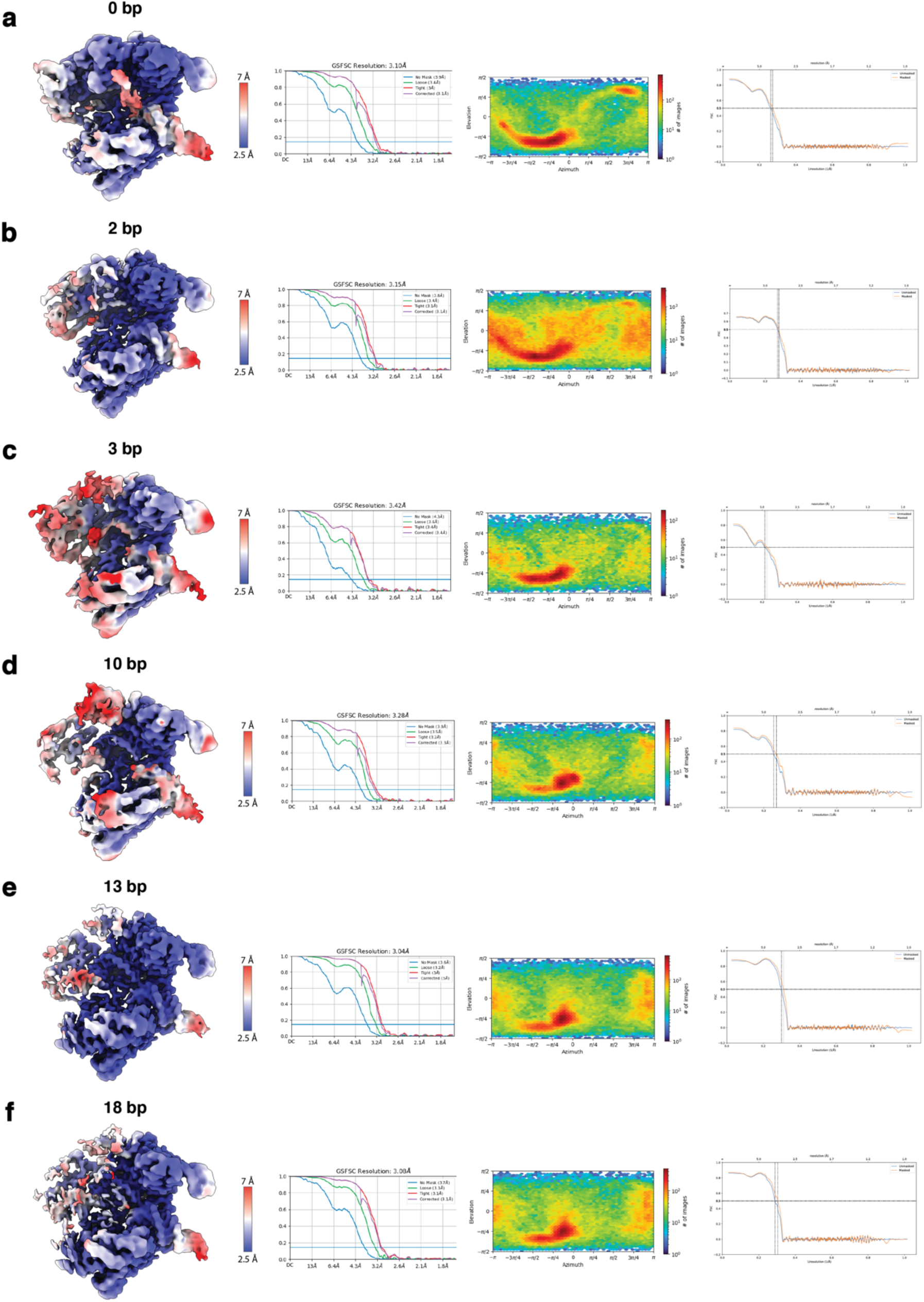
Cryo-EM data analysis of SpRY **a**, 0 bp **b**, 2 bp **c**, 3 bp **d**, 10 bp **e**, 13 bp **f**, 18 bp, and **g**, 20 bp (product state) R-loop intermediates.

**Extended Data Figure 7.**
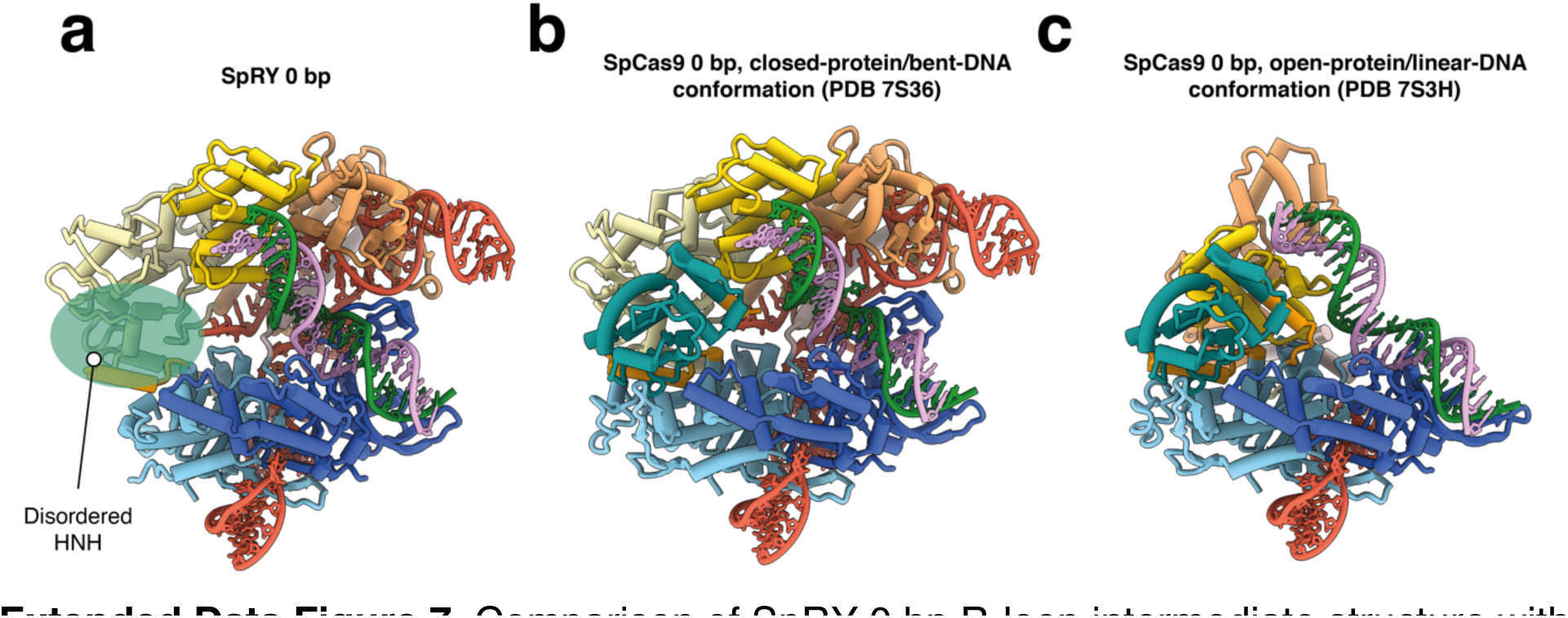
Comparison of SpRY 0 bp R-loop intermediate structure with previously determined Cas9 0 bp structures.

**Extended Data Figure 8.**
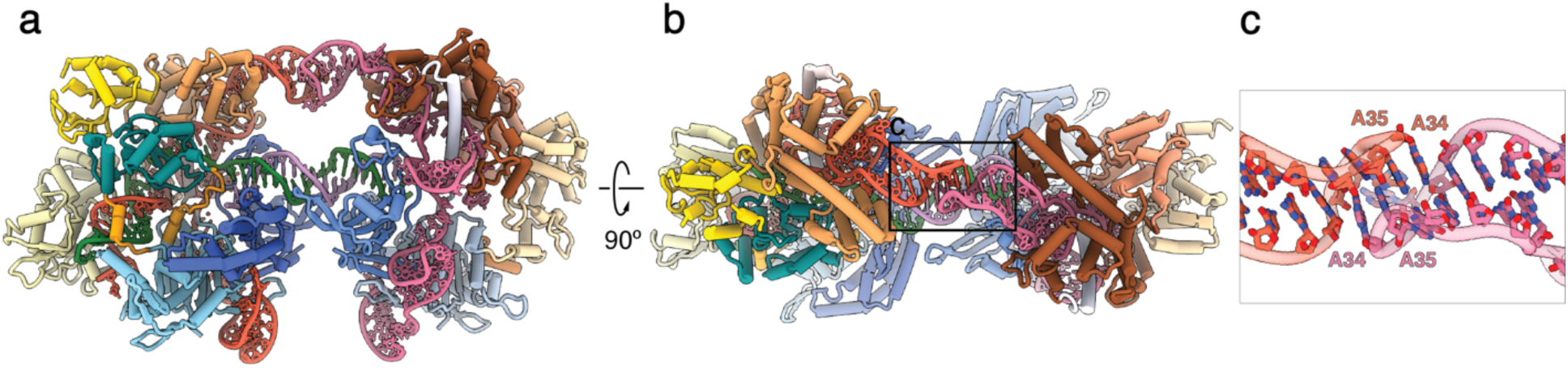
SpRYmer stem loop interaction. **a**, Overview of the SpRYmer atomic model. **b**, Top-down view of the SpRYmer atomic model. **c**, Detailed view of the stem loop kissing interaction.

**Extended data figure 9.**
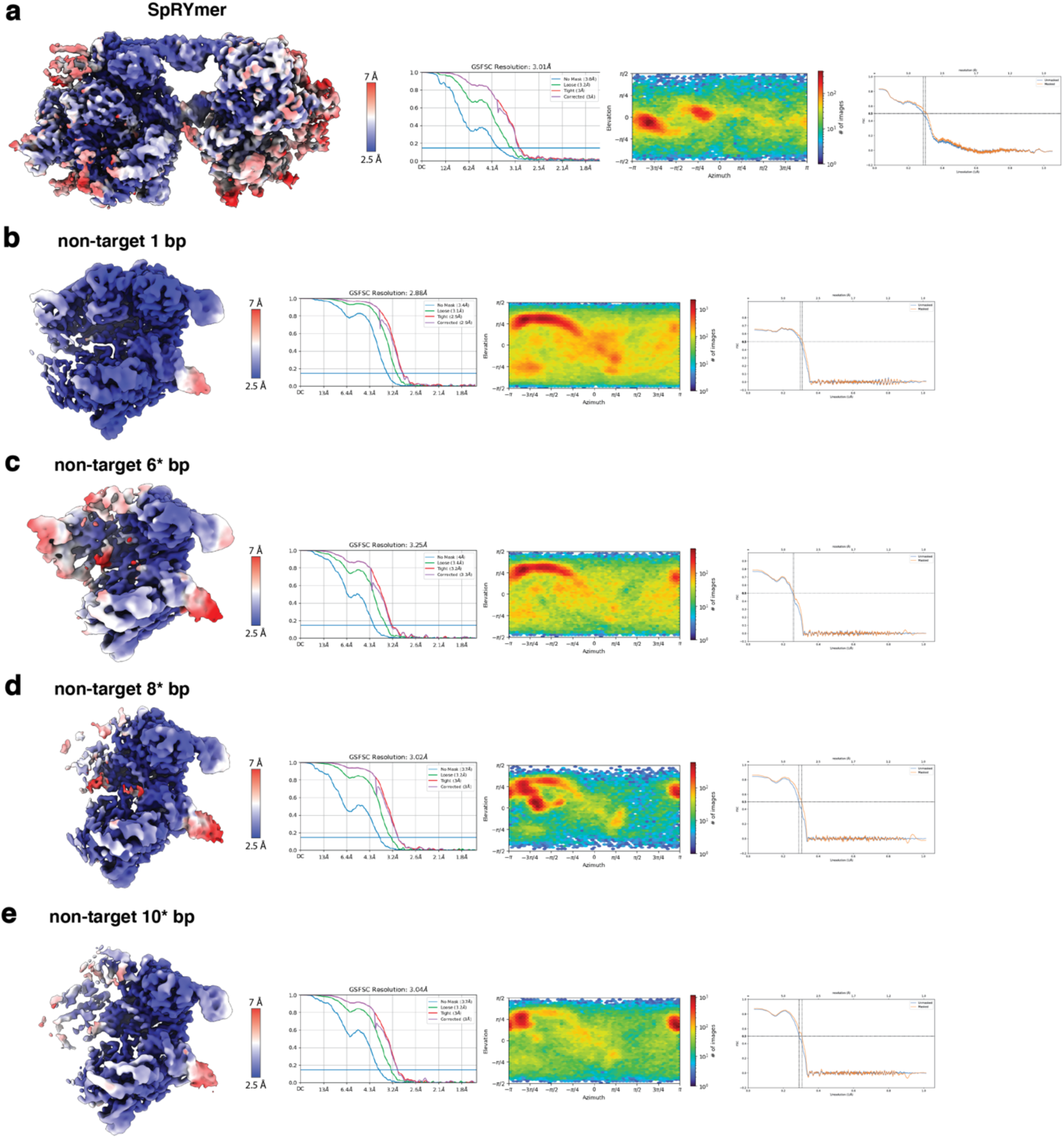
Cryo-EM data analysis of the **a**, SpRYmer **b**, SpRY bound to non-target DNA with 1 bp **c**, 6 bp **d**, 8 bp, and **e**, 10 bp of R-loop formed. *These structures contain mismatches.

**Extended Data Table 1.**
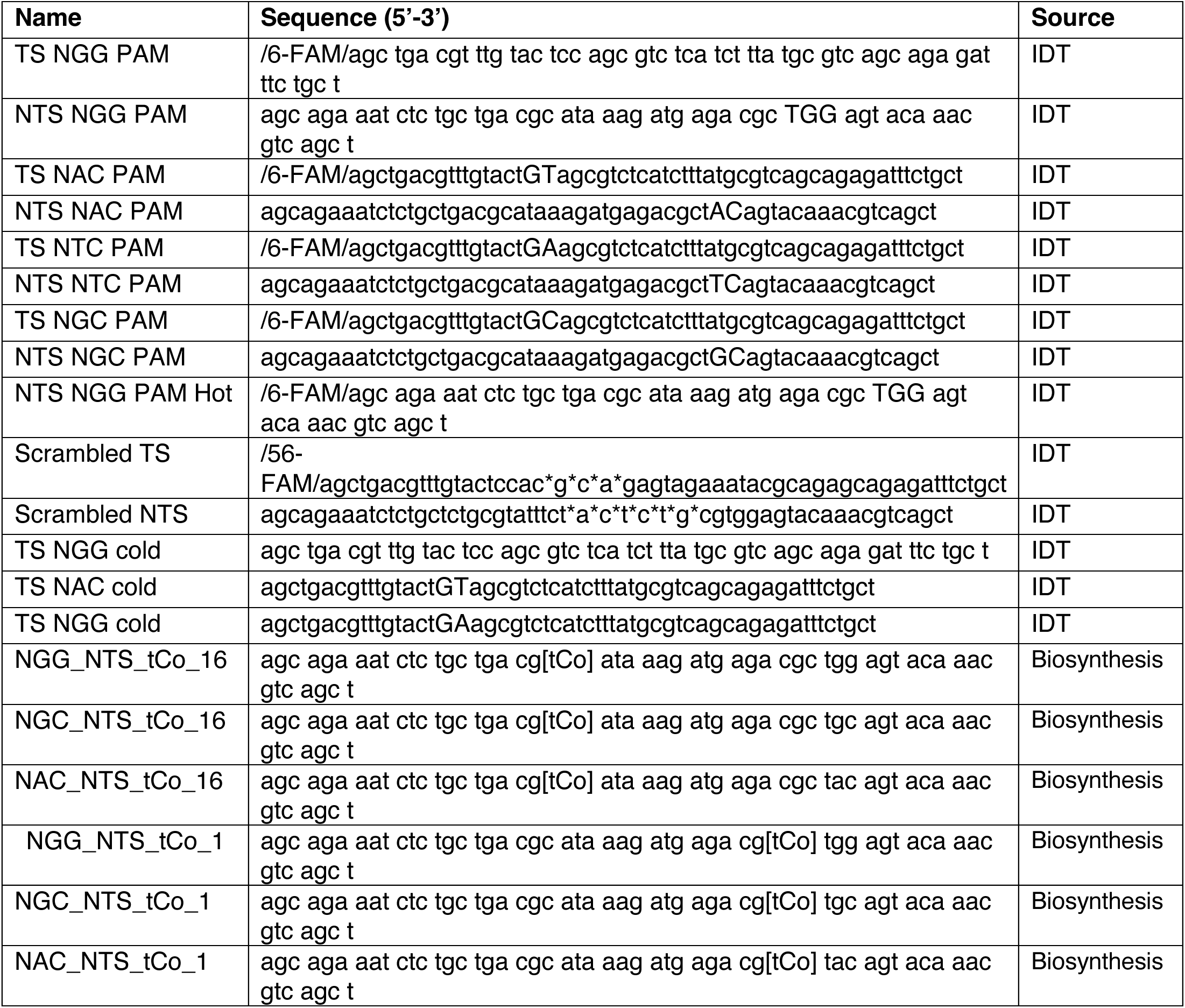
List of DNA substrates used in this study. * corresponds to phosphorotioate-modified DNA.

**Extended Data Table 2.**
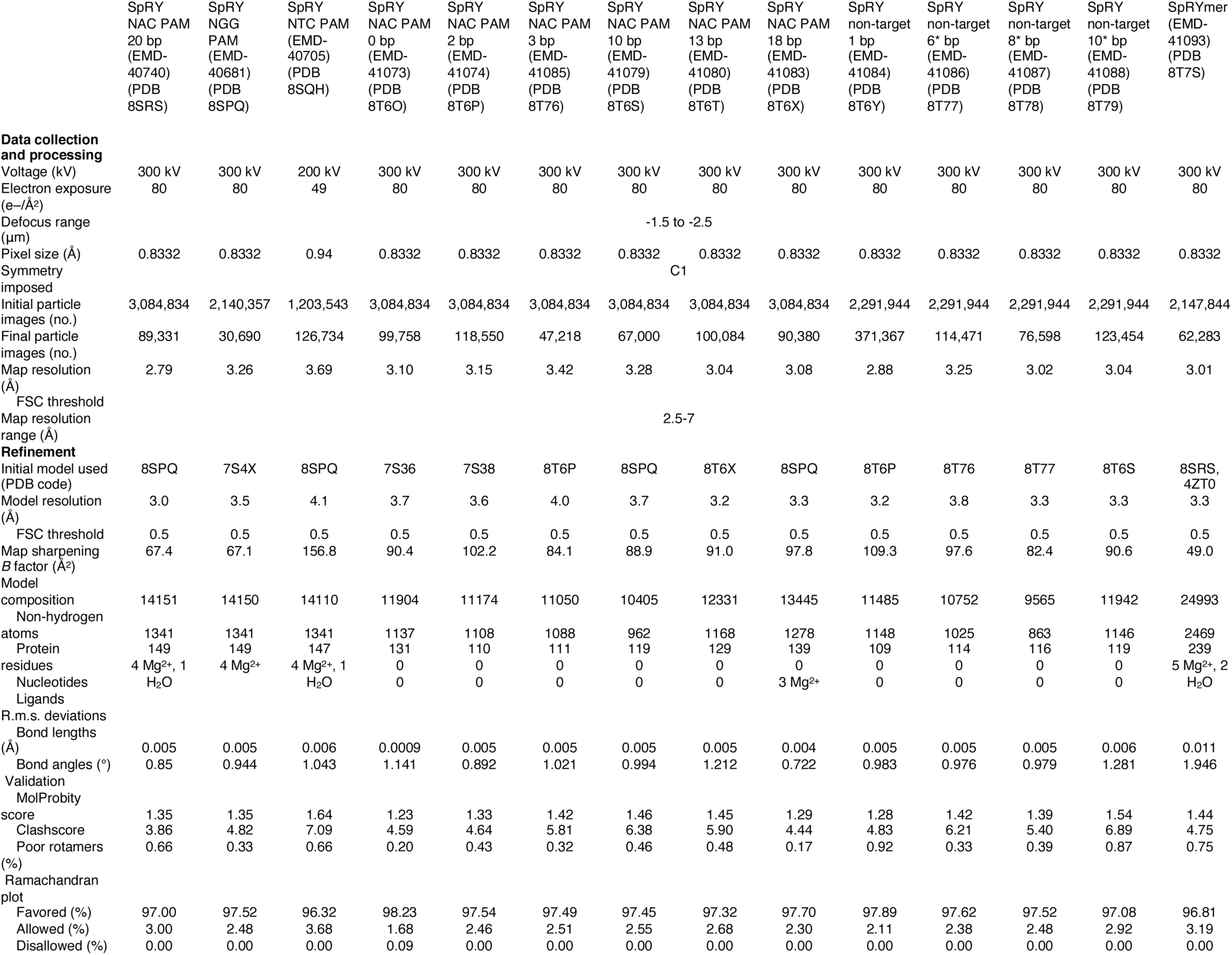
Cryo-EM data processing and modeling.

**Extended Data Table 3.** Rate constants derived from fitting the data in Extended Data Figure 3 to the model shown in **Extended Data Fig 5** are summarized. Rates for Cas9 are taken from (20). The * indicates that rates for DNA binding and dissociation were not defined by the data but were locked at these rate constants to give a K_d_ of 3 nM.

| Parameter | Best fit |  |
| --- | --- | --- |
|  | SpRY | Cas9 |
| $K_1$ | $32 \mu\text{M s}^{-1}$ | 3* |
| $K_{-1}$ | $0.094 \text{ s}^{-1}$ | |
| $K_2$ | $0.0045 \text{ s}^{-1}$ | 2.5 |
| $K_{-2}$ | $0.0003 \text{ s}^{-1}$ | 1.2 |
| $K_3$ | $20 \text{ s}^{-1}$ | 5.6 |
| $K_4$ | $0.03 \text{ s}^{-1}$ | 1,8 |
| $K_{-4}$ | 0.63 | 2.3 |
| $K_5$ | 0.54 | 4.4 |

